# Re-polyadenylation occurs predominantly on maternal mRNA degradation intermediates during mammalian oocyte-to-embryo transition

**DOI:** 10.1101/2021.08.29.458080

**Authors:** Yusheng Liu, Yiwei Zhang, Hu Nie, Zhonghua Liu, Jiaqiang Wang, Falong Lu

## Abstract

The nascent mRNA transcribed in the nucleus is cleaved and polyadenylated before it is transported to the cytoplasm for translation^1^. Polyadenylation can also occur in the cytoplasm for post-transcriptional regulation, especially in neurons, oocytes and early embryos^1,2^. Recently, we revealed transcriptome-wide maternal mRNA cytoplasmic re-polyadenylation during the mammalian oocyte-to-embryo transition (OET)^3-6^. However, the mechanism of re-polyadenylation during mammalian OET, including the sites to be re-polyadenylated and the enzymes involved, is still poorly understood. Here, by analyzing the PAIso-seq1 and PAIso-seq2 poly(A) inclusive transcriptome data during OET in mice, rats, pigs, and humans, we reveal conserved re-polyadenylation of mRNA degradation intermediates. These re-polyadenylated mRNA degradation intermediates account for over half of the polyadenylated mRNA during OET in all four species. We find that mRNA degradation intermediates for re-polyadenylation are generated through Btg4-mediated deadenylation in both mouse and human. Interestingly, the poly(A) tails on the re-polyadenylated mRNA degradation intermediates are of different lengths and contain different levels of non-A residues compared to regular polyadenylation sites, suggesting specific regulation and function of these poly(A) tails in mammalian OET. Together, our findings reveal the maternal mRNA degradation intermediates as substrates for conserved cytoplasmic dominant re-polyadenylation during mammalian OET, and uncover the mechanism of production of these mRNA degradation intermediates. These findings provide new insights into mRNA post-transcriptional regulation, and a new direction for the study of mammalian OET.

## Introduction

Most eukaryotic mRNA acquires a non-templated poly(A) tail co-transcriptionally. Pre-mRNA can be recognized by the cleavage and polyadenylation specificity factor (CPSF) at the polyadenylation signal (PAS) sequence (AAUAAA). This signal is often present 10 – 30 nt upstream of the cleavage site, which initiates cleavage and polyadenylation^1,7,8^. Therefore, a clear A-rich signal is observed about 10 – 30 nt upstream of regular polyadenylation sites^7,8^. After cleavage, the 3′-ends are polyadenylated by polyadenylate polymerase (PAP). In general, canonical PAPs catalyze all co-transcriptional polyadenylation events in the nucleus^9^.

Polyadenylation can also occur in the cytoplasm in response to inflammation, in neurons for localized translational activation, and in oocytes for translational activation of the maternal factor, important for meiosis progression^1,2,10,11^. Studies in *Xenopus* oocytes revealed that cytoplasmic polyadenylation element (CPE) is an important cis-element normally present in the 3′-UTR of mRNA that undergoes cytoplasmic polyadenylation^12,13^. This element can be recognized by cytoplasmic polyadenylation element binding protein (CPEB)^13^. CPEB1 is a bifunctional regulator that initially contributes to deadenylation by recruiting PARN deadenylase^14^. However, once phosphorylated, CPEB1 can promote cytoplasmic polyadenylation through Gld2, a non-canonical polyadenylate polymerase (ncPAP)^14^. Research into cyclin B1 in *Xenopus* oocytes revealed that both CPE and PAS are required for its cytoplasmic re-polyadenylation^14^.

Gld2 (also called Tent2) is the best known ncPAP that is able to catalyze cytoplasmic polyadenylation of CPEB-bound mRNA^15-18^. In response to developmental cues, phosphorylated CPEB1 recruits Gld2 for cytoplasmic polyadenylation, which leads to polyadenylation-induced activation of translation^14^. In *Drosophila*, Wispy, the *Drosophila* homolog of Gld2, promotes global poly(A) tail elongation during late oogenesis^19-22^. There are 11 ncPAPs, which are also called terminal nucleotidyltransferases (TENTs), in the mammalian genome^15,16^. Besides Gld2/Tent2, Tent4 and Tent5 family enzymes are also able to catalyze cytoplasmic polyadenylation for mRNA stabilization and translation activation^15^. In particular, the Tent4 family enzymes are capable of mix-tailing the mRNA with non-A residues, particularly G residues, which can impede the deadenylase complex^23^. We recently found extensive cytoplasmic re-polyadenylation events during OET in mice, rats, pigs and humans^3-6^. Although Gld2 family enzymes are important for cytoplasmic re-polyadenylation in *Xenopus* and *Drosophila* oocytes as well as mammalian somatic cells^18-22,24^, most likely Gld2/Tent2 is not responsible for cytoplasmic re-polyadenylation during mammalian OET, as *Tent2* knockout (KO) female mice retain normal fertility^25^.

Although the biochemical properties of pre-mRNA co-transcriptional polyadenylation have been elucidated in the nucleus, the substrates for cytoplasmic re-polyadenylation of maternal RNA in mammalian oocytes and embryos are poorly studied. In this study, we employ transcriptome-wide poly(A) inclusive PAIso-seq1 and PAIso-seq2 data of mammalian OET in mouse, rat, pig and human to study the re-polyadenylation sites during mammalian OET. Surprisingly, we reveal that re-polyadenylation predominantly occurs on mRNA degradation intermediates which are conserved and represent more than half of the poly(A)+ mRNA in all four species. The degradation intermediate substrates are generated through Btg4 dependent global deadenylation in mouse and human. Interestingly, the poly(A) tails on these degradation intermediates are of different length and contain non-A residues.

## Results

### Unique nucleotide composition around polyadenylation sites during mouse oocyte-to-embryo transition

mRNA is regulated post-transcriptionally during mouse OET before ZGA, because no new transcription happens during this process. When analyzing the alternative polyadenylation sites of polyadenylated mRNA during mouse OET (including GV: germinal vesicle, MI: metaphase I, and MII: metaphase II stage oocytes as well as 1C: 1-cell, 2C: 2-cell, and 4C: 4-cell stage embryos), we notice that there are many mRNA reads in 1C embryos that cannot be assigned to the polyadenylation sites called in GV oocytes. This suggests global change of mRNA polyadenylation sites during mouse OET. Therefore, we first looked into nucleotide composition around polyadenylation sites during mouse OET. The nucleotide composition around polyadenylation sites in the GV, MI and 4C PAIso-seq1 data showed characteristic features of commonly known polyadenylation sites in mammals (Fig. 1a), including apparent enrichment of A nucleotides 10 – 30 nt upstream of the polyadenylation sites which is the feature of the common polyadenylation signal (PAS) motif AAUAAA^7,8,26^. Surprisingly, nucleotide composition around the polyadenylation sites of 1C data differs dramatically from the GV data as measured by PAIso-seq1 (Fig. 1a). 1C shows reduced signals of common features, while the nucleotide composition of the MII and 2C data falls between the 1C and GV stages (Fig. 1a). To further confirm this observation, we looked into a PAIso-seq2 dataset which captures mRNA 3′-ends through direct ligation of 3′ adaptors^27^. A similar phenomenon is observed in the PAIso-seq2 dataset (Fig. 1b), confirming the reliability and reproducibility of our observations. Together, the results from both the PAIso-seq1 and PAIso-seq2 datasets reveal that polyadenylation sites around the 1C stage in mouse are of very different surrounding nucleotide composition, suggesting global changes to polyadenylation sites during mouse OET.

**Fig. 1.**
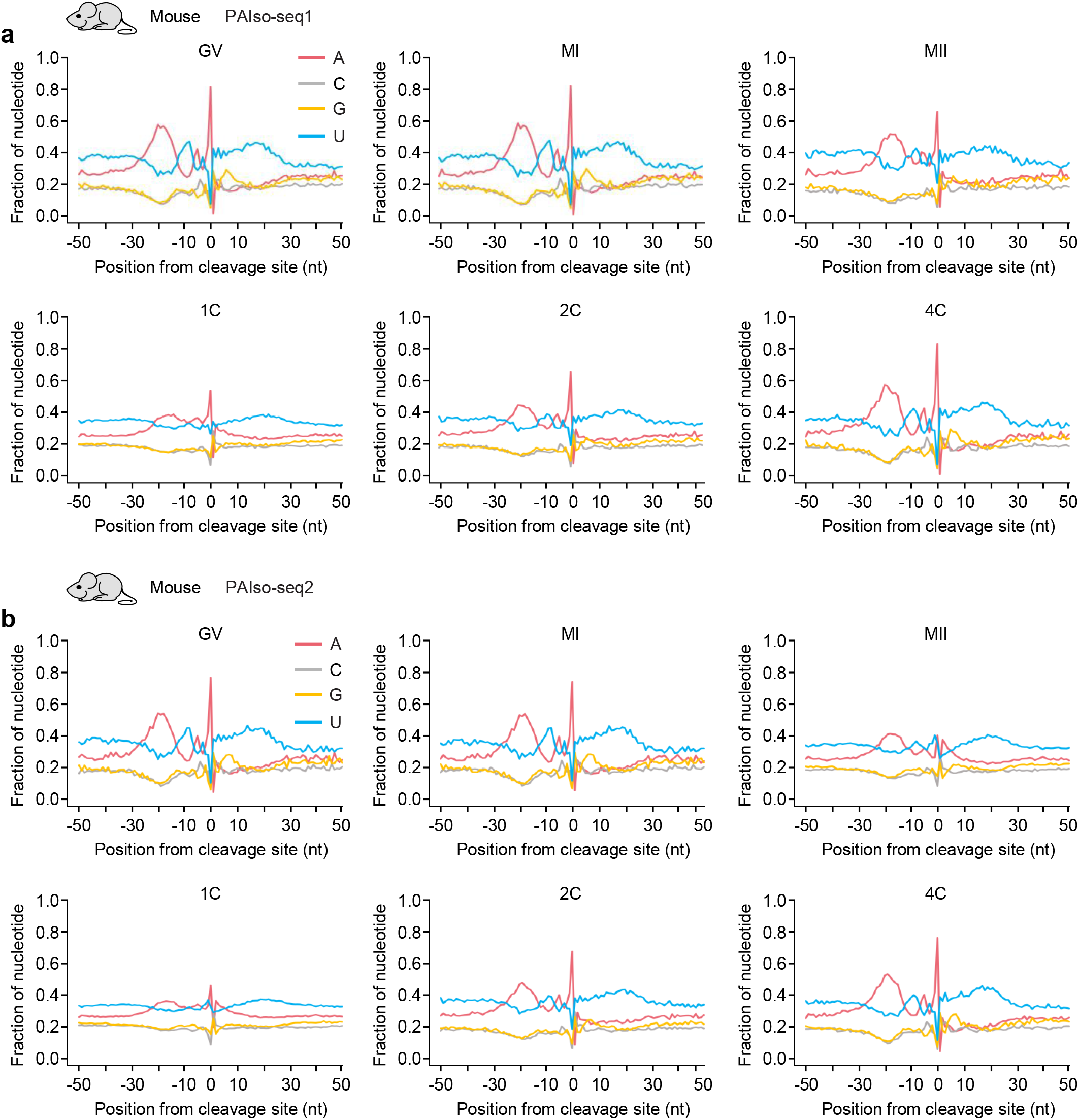
Nucleotide composition around the polyadenylation sites during mouse oocyte-to-embryo transition. Profile of nucleotide frequencies of the genomic sequences in the ± 50 nt vicinity of the last base before mRNA poly(A) tails sequenced by PAIso-seq1 (**a**) or PAIso-seq2 (**b**) in samples from different stages during mouse OET. GV, germ-vesicle oocyte; MI, metaphase I oocyte; MII, metaphase II oocyte; 1C, 1-cell embryo; 2C, 2-cell embryo; 4C, 4-cell embryo. Transcripts with a poly(A) tail of at least 1 nt are included in the analysis.

### Re-polyadenylated mRNA degradation intermediates dominate polyadenylated mRNA during mouse oocyte-to-embryo transition

Next, we used the genome browser to look into the mRNA 3′-ends sequenced by PAIso-seq1. In the GV stage, the majority of polyadenylation events occur within a small window around the known polyadenylation sites, for example, almost all of the mRNA of *Plat* and *Tle6* polyadenylate at the annotated 3′-ends (Fig. 2a). However, from the MI stage, we could see that part of the polyadenylation sites deviated from the annotated 3′-end toward the 5′ direction for *Plat* mRNA, which became more evident in MII and 1C stages along with development (Fig. 2a). This resulted in a continuous distribution of polyadenylation sites upstream of the original polyadenylation sites (Fig. 2a). As there is no new transcription, these observations reveal that a large amount of mRNA become partially degraded in the 3′-ends, which can then be re-polyadenylated to generate re-polyadenylated mRNA degradation intermediates during mouse OET (Fig. 2b). We called the poly(A) tails added at the original polyadenylation sites classical poly(A) tails, and those added 5′ away from the original polyadenylation sites non-classical poly(A) tails (Fig. 2b).

**Fig. 2.**
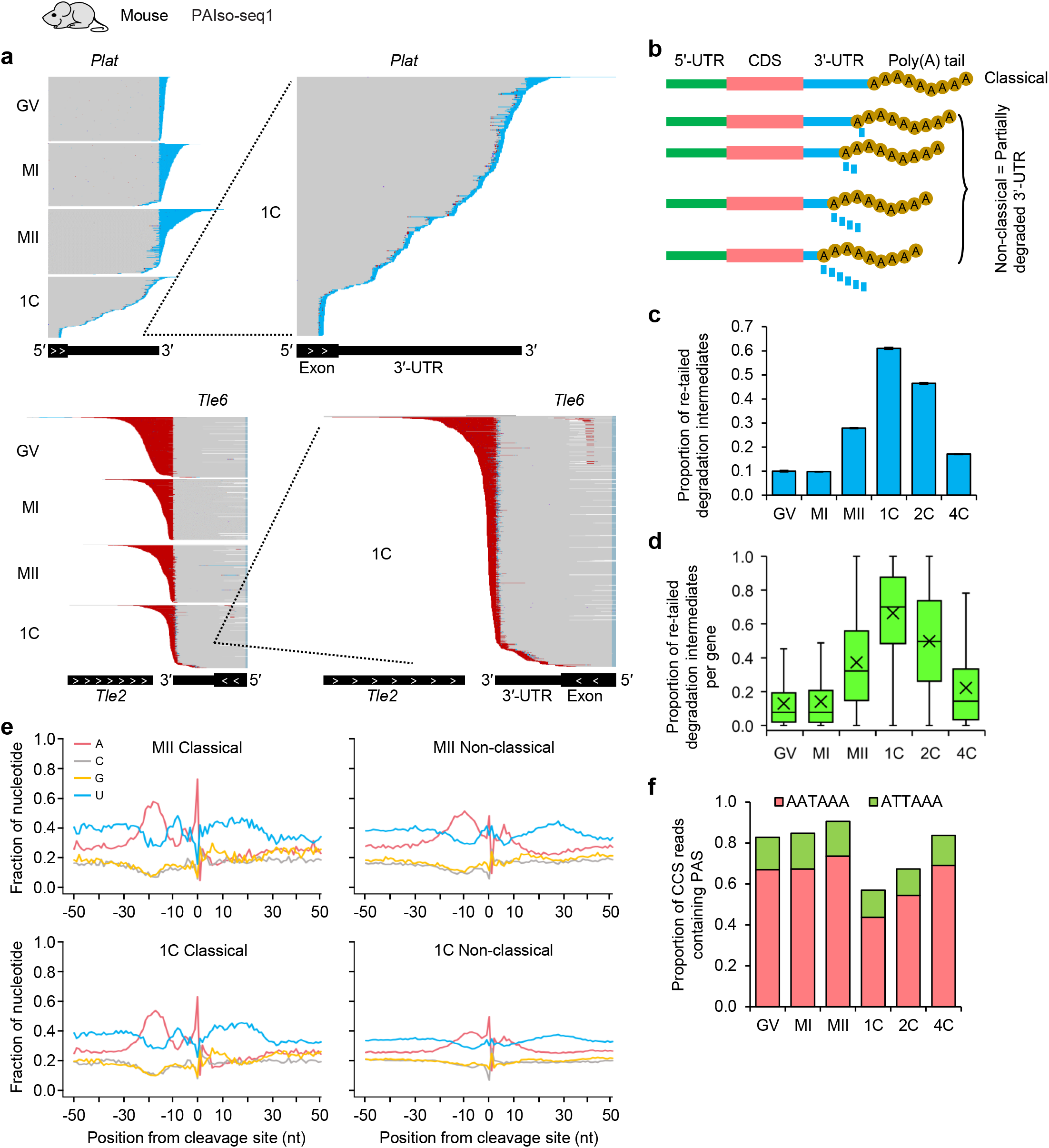
Re-polyadenylated mRNA degradation intermediates dominate polyadenylated mRNA during mouse oocyte-to-embryo transition. **a**, Integrative genomics viewer (IGV) tracks showing PAIso-seq1 CCS reads of *Plat* and *Tle6* near the annotated polyadenylation sites in GV, MI, MII and 1C mouse samples sequenced by PAIso-seq1. The tracks of 1C samples are magnified at the right. The sequences that match the reference genome are shown in grey; the A (or T in the reverse direction) residues of the poly(A) tail are shown in cyan (or red). **b**, Illustration of transcripts with classical or non-classical polyadenylation sites. **c**, Overall proportion of the re-polyadenylated mRNA degradation intermediates for each of the 6 stage mouse samples sequenced by PAIso-seq1. **d**, Box plot of the proportion of the re-polyadenylated mRNA degradation intermediates for each gene in each of the 6 stages of mouse samples sequenced by PAIso-seq1. Genes (n = 4,889) with at least 10 transcripts with poly(A) tail of at least 1 nt in each of the 6 stages are included in the analysis. The “×” indicates the mean value, the black horizontal bars show the median value, and the top and bottom of the box represent the value of 25^th^ and 75^th^ percentile, respectively. **e**, Profile of nucleotide frequencies of the genomic sequences in the ± 50 nt vicinity of the last base before mRNA poly(A) tails for the combined transcripts with poly(A) tails from classical (left) and non-classical (right) polyadenylation sites in MII (top) and 1C (bottom) mouse samples sequenced by PAIso-seq1. **f**, Overall proportion of mRNA transcripts with canonical polyadenylation signal (PAS) in each of the 6 stage mouse samples sequenced by PAIso-seq1. Error bars indicate the SEM from two replicates (n=2). Transcripts with a poly(A) tail of at least 1 nt are included in the analysis.

We quantified the non-classical poly(A) tails in each of the stages of the PAIso-seq1 data. We found that non-classical poly(A) tails were at baseline levels in the GV, MI and 4C stage samples, and peaked at the 1C stage, representing more than 60% of the poly(A) tails (Fig. 2c). At the individual gene level, we observed similar pattern of non-classical poly(A) tails during mouse OET (Fig. 2d). This increase of the non-classical poly(A) tails coincides with the observed reduction of the common nucleotide composition feature around polyadenylation sites (Fig. 1). Therefore, it is very likely that the appearance of non-classical poly(A) tails explains the reduction of the common nucleotide composition feature around polyadenylation sites. Indeed, in differentiating the classical and the non-classical poly(A) tails, the classical poly(A) tails shared a common feature of nucleotide composition around the polyadenylation sites, whereas the non-classical poly(A) tails showed a reduction in this signal in both MII and 1C PAIso-seq1 data, including the A rich signal corresponding to the PAS (Fig. 2e). As there is no new transcription during this period, we reason that the reduction of this nucleotide signal might be due to the degradation of the maternal mRNA transcripts along the 3′UTRs, and some of PASs has degraded during this process. These mRNA degradation intermediates without PAS can still be re-polyadenylated. Indeed, a large fraction of the polyadenylated mRNA transcripts are devoid of canonical PAS and its most common variant in mouse 1C embryos compared to GV and 4C stages (Fig. 2f). Moreover, the above observations in the PAIso-seq1 dataset were faithfully recaptured in the PAIso-seq2 dataset, which further confirmed our observations (Extended Data Fig. 1).

As there is no new transcription between GV and 1C stages, the unique feature of polyadenylation sites seen in the 1C stage must be a result of post-transcriptional regulation of the maternal mRNA. Our studies have recently revealed the existence of global deadenylation and re-polyadenylation events during mouse OET^4,5^. However, the substrate for the global re-polyadenylation is not clear. Therefore, the above results reveal that the mRNA partially-degraded intermediates (called mRNA degradation intermediates hereafter) are the main substrate for re-polyadenylation during mouse OET, which results in re-polyadenylated mRNA degradation intermediates as the dominant pool of polyadenylated mRNA in mouse 1C embryos.

### Maternal mRNA deadenylation is required for the generation of re-polyadenylated mRNA degradation intermediates

The above results suggest that the non-classical poly(A) tails are added to mRNA degradation intermediates through re-polyadenylation. As deadenylation precedes degradation, we reason that if we inhibit the deadenylation process, we expect reduced levels of mRNA degradation intermediates, leading to a reduction of the non-classical poly(A) tails. We and others previously demonstrated that Btg4 regulates global mRNA deadenylation during mouse oocyte maturation^4,5,28-30^. Therefore, we look into the level of non-classical poly(A) tails after *Btg4* KD. Consistent with our predictions, we found that the level of total non-classical poly(A) tails decreased significantly in *Btg4* KD MII oocytes (Fig. 3a). This decrease was also evident at the individual gene level (Fig. 3b). Alternatively, we tried to block mRNA deadenylation by treating oocytes with adenosine monophosphate (AMP), which inhibits the general active deadenylation of poly(A) tails, by potent inhibition of the major de-adenylating enzymes Cnot6l and Cnot7^31^. Similar to what we observed in *Btg4* KD, we saw a significant decrease in non-classical poly(A) tails for the total mRNA pool and for individual genes after AMP treatment (Fig. 3c, d). Together, these data demonstrate that the high level of non-classical tails seen during mouse OET are generated through re-polyadenylation of the mRNA degradation intermediates, which are produced by Btg4 mediated deadenylation followed by 3′-UTR partial degradation through 3′-5′ exonucleases.

**Fig. 3.**
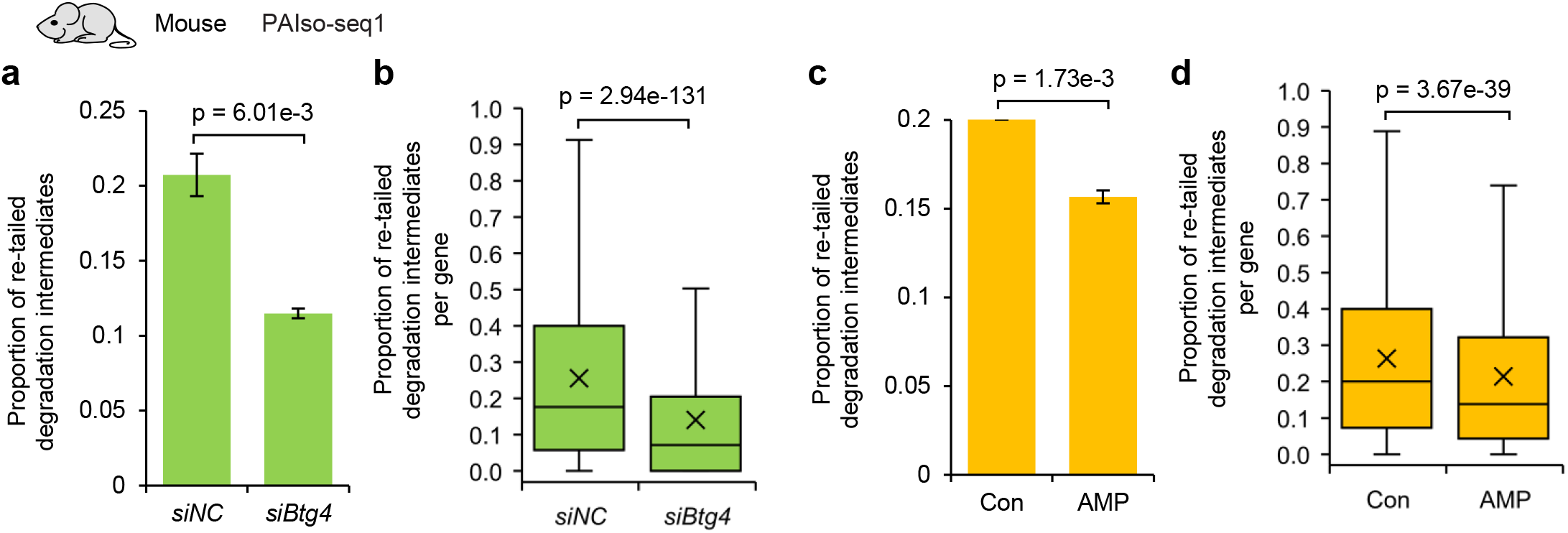
Maternal mRNA deadenylation is required for the generation of re-polyadenylated mRNA degradation intermediates. **a**, Overall proportion of the re-polyadenylated mRNA degradation intermediates in *siNC* and *siBtg4* mouse MII oocytes sequenced by PAIso-seq1. **b**, Box plot of the proportion of the re-polyadenylated mRNA degradation intermediates for each gene in *siNC* and *siBtg4* mouse MII oocytes sequenced by PAIso-seq1. Genes (n = 3,154) with at least 10 transcripts with a poly(A) tail of at least 1 nt in both samples are included in the analysis. **c**, Overall proportion of the re-polyadenylated mRNA degradation intermediates in mouse MII oocytes treated with or without AMP sequenced by PAIso-seq1. **d**, Box plot of the proportion of the re-polyadenylated mRNA degradation intermediates for each gene in mouse MII oocytes treated with or without AMP sequenced by PAIso-seq1. Genes (n = 7,048) with at least 10 transcripts with a poly(A) tail of at least 1 nt in both samples are included in the analysis. All the *p* values are calculated by Student’s *t*-test. Error bars indicate the SEM from two replicates (n=2). Transcripts with a poly(A) tail of at least 1 nt are included in the analysis.

### Classical and non-classical poly(A) tails are different

The non-classical poly(A) tails are highly abundant during mouse OET. We asked whether the classical and non-classical poly(A) tails are similar or unique in length or non-A residue content. We found that overall, the classical poly(A) tails are longer than the non-classical poly(A) tails in both MII oocytes and 1C embryos as measured by both PAIso-seq1 and PAIso-seq2 (Fig. 4a, d). In addition, we observed significantly higher levels of non-A residues within the non-classical poly(A) tails compared to the non-classical poly(A) tails in general in both MII oocytes and in 1C embryos as measured by both PAIso-seq1 and PAIso-seq2 (Fig. 4b, c, e, f). The difference in the length and the non-A residue level between classical and non-classical poly(A) tails is likely caused by their origin and mechanism of synthesis. The non-classical poly(A) tails are generated through new re-polyadenylation on the mRNA degradation intermediates, while the majority of the classical poly(A) tails are inherited from the stable maternal mRNA generated co-transcriptionally. The different features of the non-classical poly(A) tails from the classical poly(A) tails suggest that these newly synthesized non-classical poly(A) tails may be of important regulatory function during mouse OET.

**Fig. 4.**
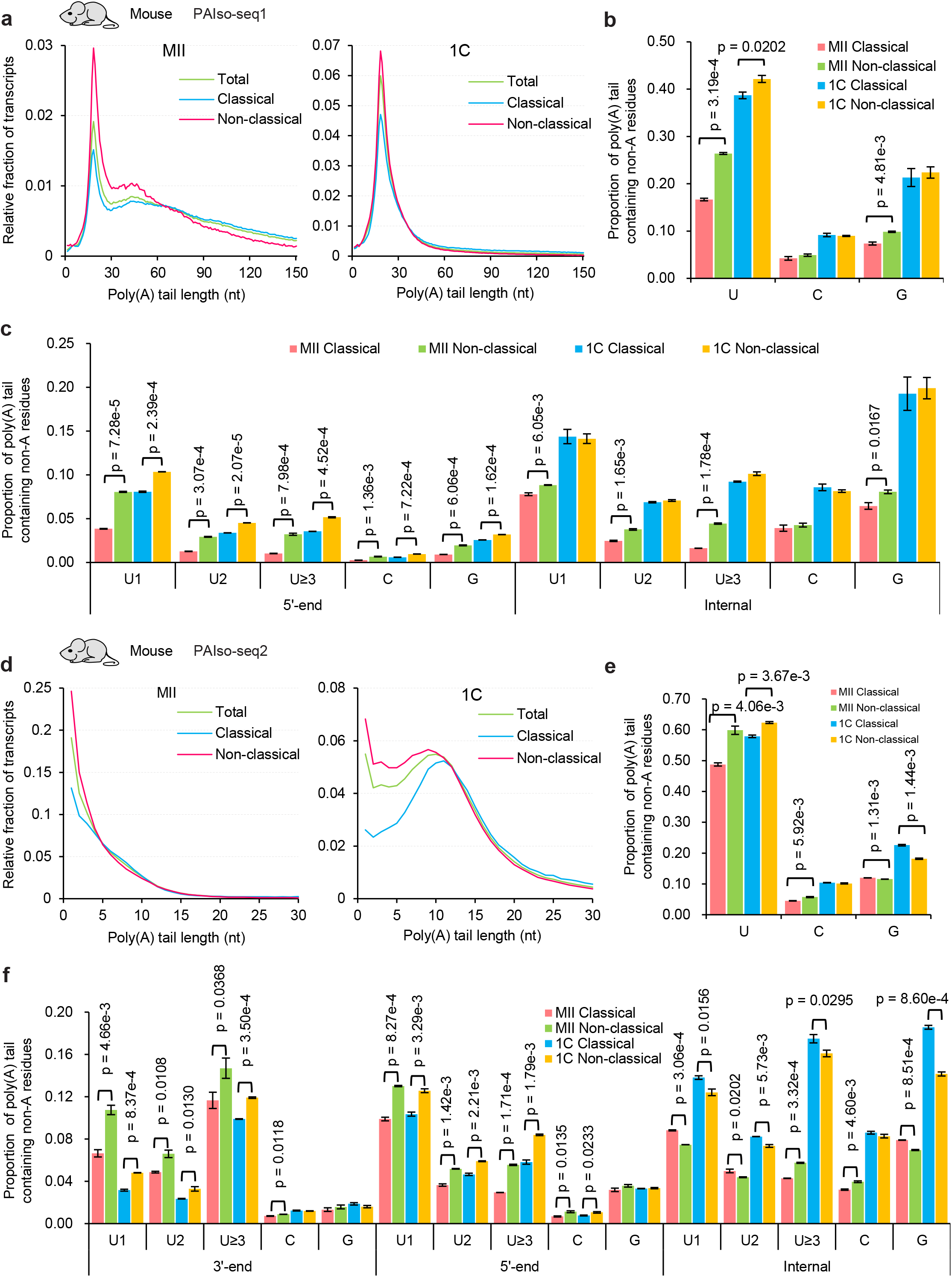
Comparison of poly(A) tails at classical and non-classical polyadenylation sites. **a, d**, Histogram of poly(A) tail lengths of all transcripts, transcripts with poly(A) tails from classical and non-classical polyadenylation sites in MII (left) and 1C (right) mouse samples measured by PAIso-seq1 (**a**) or PAIso-seq2 (**d**). Histograms (bin size = 1 nt) are normalized by the total counts of mRNA reads with a poly(A) tail of at least 1 nt. **b, e**, Overall proportion of transcripts containing U, C, or G residues in poly(A) tails from classical and non-classical polyadenylation sites in MII and 1C mouse samples measured by PAIso-seq1 (**b**) or PAIso-seq2 (**e**). **c, f**, Proportion of transcripts with U, C, or G residues at the indicated positions (5′-end, internal, and 3′-end if available) of poly(A) tails from classical and non-classical polyadenylation sites in MII and 1C mouse samples measured by PAIso-seq1 (**c**) or PAIso-seq2 (**f**). The U residues was further divided according to the length of the longest consecutive U (1, 2, and ≥3). All the *p* values are calculated by Student’s *t*-test. Error bars indicate the SEM from two replicates (n=2). Transcripts with a poly(A) tail of at least 1 nt are included in the analysis.

### Similar nucleotide composition around polyadenylation sites in all the examined somatic cells from mouse, rat, pig and human

As the non-classical poly(A) tails represent more than half of the poly(A) tails in 1C embryos, we asked whether the non-classical poly(A) tails are also present in somatic cells. We investigated somatic cell poly(A) tail datasets by PAIso-seq1, PAIso-seq2 and FLAM-seq from mouse, rat, pig and human, including mouse liver^32^, mouse 3T3 cells^27^, mouse embryonic stem (ES) cells^27^, rat liver, pig liver, and human HeLa cells^33^. We found that the nucleotide composition around the polyadenylation sites of these somatic cells matched characteristic features of commonly known polyadenylation sites (Fig. 5a-g). In addition, the nucleotide composition around the polyadenylation sites of 12 additional mouse tissue samples, including brown adipose tissue (BAT), brain, epididymis, heart, intestine, kidney, muscle, ovary, oviduct, lung, testis and uterus, matched characteristic features of commonly known polyadenylation sites (Extended Data Fig. 2). Therefore, these data suggest that the non-classical poly(A) tails are unique features to the mouse OET process.

**Fig. 5.**
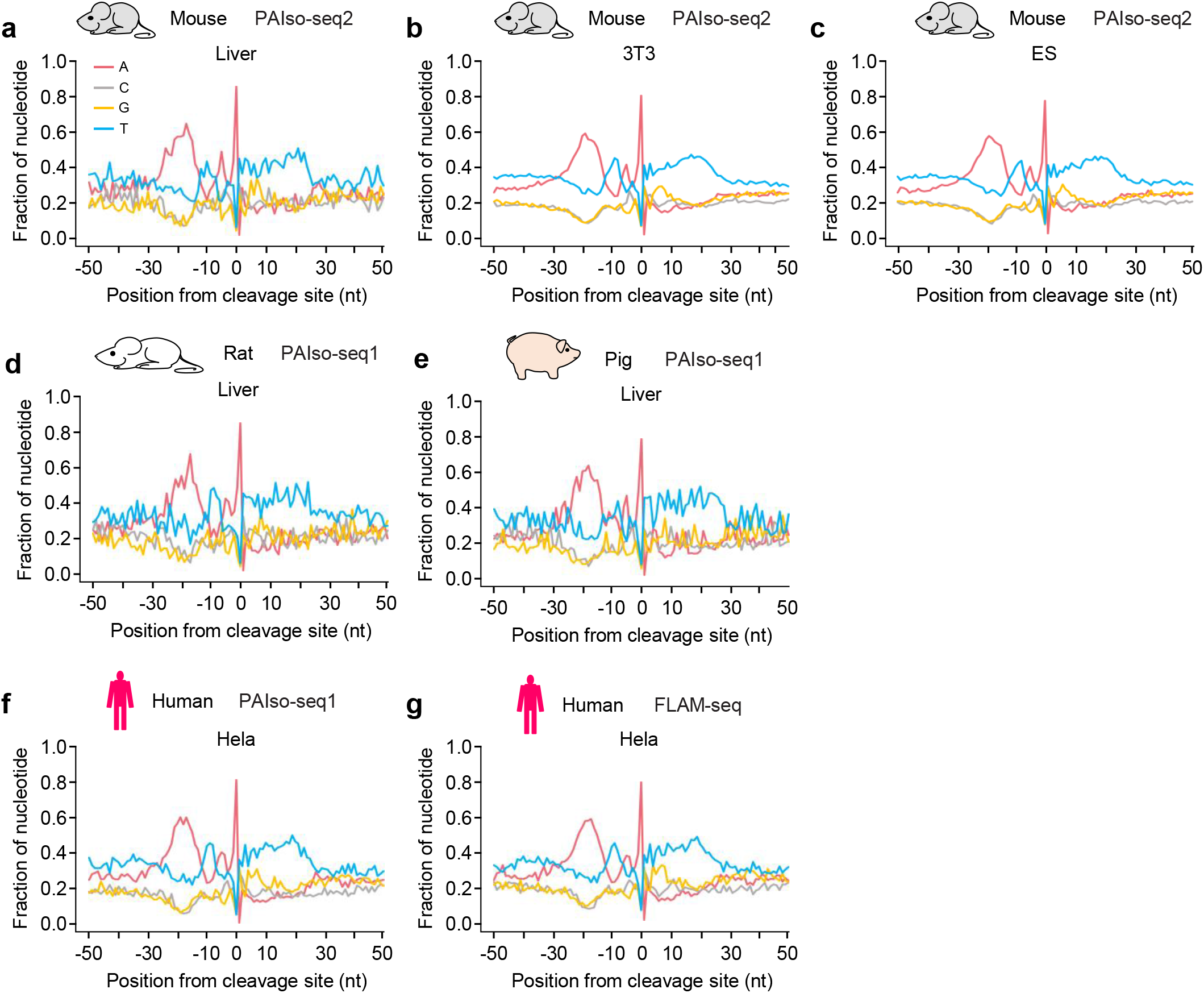
Similar nucleotide composition around the polyadenylation sites in somatic cells from mouse, rat, pig and human. Profile of nucleotide frequencies of the genomic sequences in the ± 50 nt vicinity of the last base before mRNA poly(A) tails sequenced by PAIso-seq1 in mouse liver (**a**), mouse 3T3 cells (**b**) and mouse embryonic stem cells (**c**) by PAIso-seq2, rat liver (**d**), pig liver (**e**) and human HeLa cells (**f**) by PAIso-seq1, as well as human HeLa cells by FLAM-seq (**g**).

### Conserved domination of polyadenylated mRNA by re-polyadenylated mRNA degradation intermediates during oocyte-to-embryo transition of rat, pig and human

Next, we asked whether the non-classical poly(A) tails are unique features of mouse OET or conserved features across mammalian OET. Therefore, we looked in the PAIso-seq2 data of different stage OET samples in rat, pig and human^3,6^. We found that rat exhibits the same nucleotide composition features around polyadenylation sites as mouse in GV, MII, 1C, and 2C samples (Fig. 6a-c). For pig samples, we found that the nucleotide composition features around polyadenylation sites are similar to mouse in the GV, MII and 1C stages, while the pig 2C stage sample resembled the 1C stage in pig (Fig. 6d-f). For human samples, we found that the nucleotide composition features around polyadenylation sites is similar to mouse in the GV, MII and 1C stages, while the human 2C and 4C stage samples exhibit even further reduction of the characteristic features of commonly known polyadenylation sites and increase of the non-classical poly(A) tails compared to the human 1C sample (Fig. 6g-i).

**Fig. 6.**
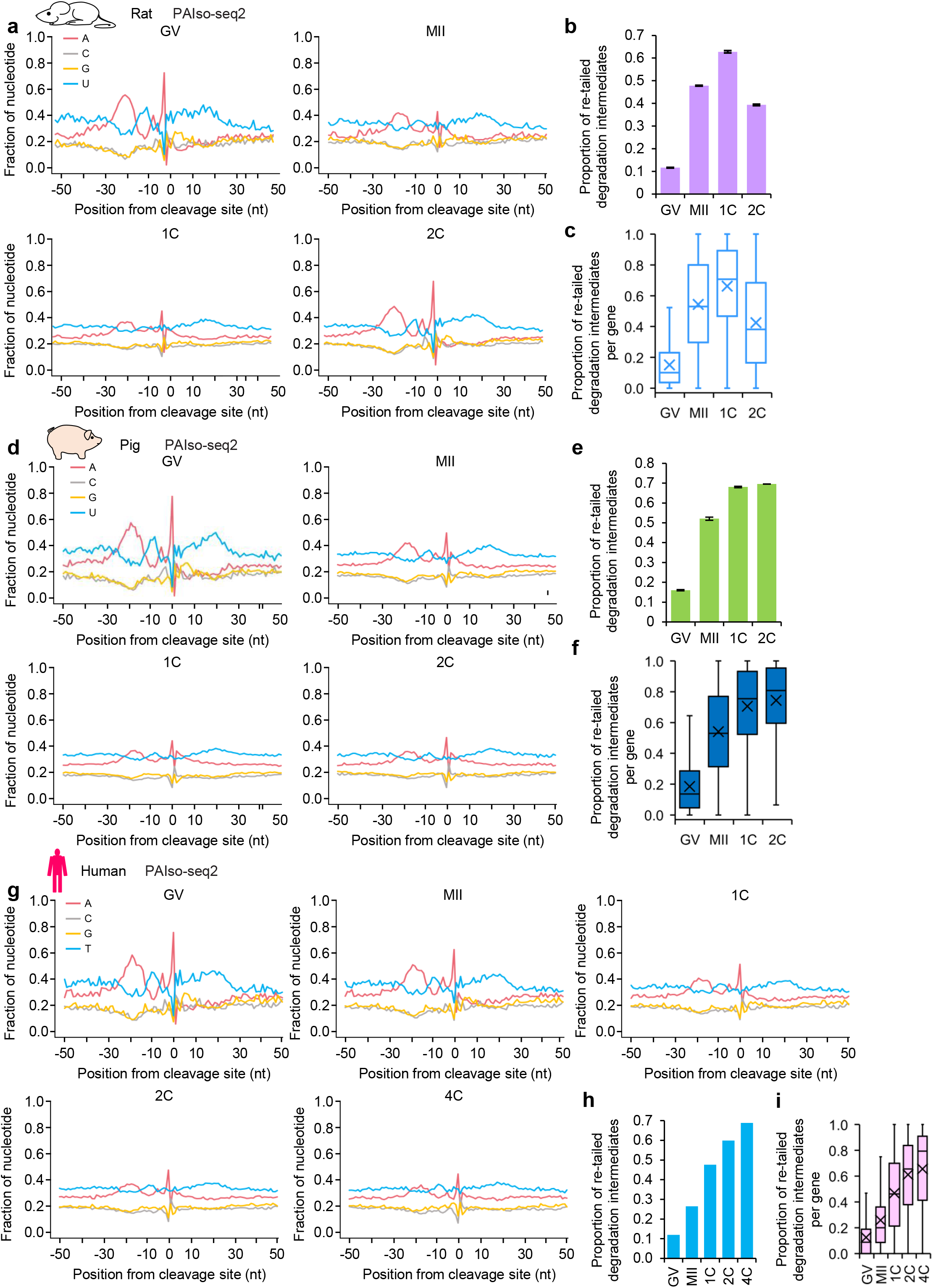
Re-polyadenylated mRNA degradation intermediates dominate polyadenylated mRNA during oocyte-to-embryo transition of rat, pig, and human. **a, d**, Profile of nucleotide frequencies of the genomic sequences in the ± 50 nt vicinity of the last base before mRNA poly(A) tails sequenced by PAIso-seq2 in GV, MII, 1C and 2C stage samples in rat (**a**) and pig (**d**). **b, e**, Overall proportion of the re-polyadenylated mRNA degradation intermediates for each of the 4 stage PAIso-seq2 samples in rat (**b**) and pig (**e**). **c, f**, Box plot of the proportion of the re-polyadenylated mRNA degradation intermediates for each gene in each of the 4 stages PAIso-seq2 samples in rat (**c**) and pig (**f**). Genes (n = 1,435 for rat samples, n = 2,869 for pig samples) with at least 10 transcripts with a poly(A) tail of at least 1 nt in each of the 4 stages are included in the analysis. **g**, Profile of nucleotide frequencies of the genomic sequences in the ± 50 nt vicinity of the last base before mRNA poly(A) tails sequenced by PAIso-seq2 in GV, MII, 1C, 2C and 4C stage samples in human. **h**, Overall proportion of the re-polyadenylated mRNA degradation intermediates for each of the 5 stages of human PAIso-seq2 samples. **i**, Box plot of the proportion of the re-polyadenylated mRNA degradation intermediates for each gene in each of the 5 stages human PAIso-seq2 samples. Genes (n = 82) with at least 10 transcripts with poly(A) tail of at least 1 nt in each of the 5 stages are included in the analysis. Error bars indicate the SEM from two replicates (n=2). Transcripts with a poly(A) tail of at least 1 nt are included in the analysis. For all the box plots, the “×” indicates the mean value, the black horizontal bars show the median value, and the top and bottom of the box represent the value of 25^th^ and 75^th^ percentile, respectively.

For pig and human, the PAIso-seq1 datasets also showed that the non-classical poly(A) tails became dominant after fertilization (Extended Data Fig. 3, Extended Data Fig. 4a-c). As the human *BTG4* KD 1C PAIso-seq1 data is available^3^, we asked whether BTG4 mediated deadenylation is required for the generation of the non-classical poly(A) tails in human. Indeed, our results showed that the level of non-classical poly(A) tails decreased in human *BTG4* KD 1C embryos (Extended Data Fig. 4d-e), confirming the conserved role for BTG4 in the generation of mRNA degradation intermediates as the substrates for re-polyadenylation to generate mRNA with non-classical poly(A) tails. In human the dominant non-classical poly(A) tails range from the 1C to 4C stage, and begin to decrease in the 8C stage (Extended Data Fig. 4), while in pig the dominant non-classical poly(A) tails range from the 1C to 8C stage, and begin to decrease in the MO stage (Extended Data Fig. 3). In contrast, in mouse the decrease occurred at the 2C stage (Fig. 2c). The different timing of the decrease of non-classical poly(A) tails coincides with the ZGA timing of these species^34^, further confirming that non-classical poly(A) tails are added to the maternal mRNA. Therefore, these results reveal that the pattern of non-classical poly(A) tails added to the mRNA degradation intermediates during OET is conserved across mouse, rat, pig and human. This is consistent with our recent findings that high levels of non-A residues are incorporated into maternal mRNA during the re-polyadenylation event during mouse and human OET^3,5^. Together, our findings reveal that cytoplasmic re-polyadenylation of mRNA degradation intermediates with non-A residues are common conserved features of mammalian OET, which is essential for mammalian reproduction.

## Discussion

Post-transcriptional regulation of maternal mRNA stored in oocytes is the primary mechanism for regulating the mammalian OET before ZGA^1,35-39^. Cytoplasmic polyadenylation is one of the most important post-transcriptional regulatory mechanisms for temporal control of this process^1,2,35,36,38,39^. In this study, we reveal that cytoplasmic polyadenylation mainly happens on mRNA degradation intermediates during mammalian OET in mice, rats, pigs, and humans (Fig. 7). We demonstrate that Btg4-dependent mRNA deadenylation is required for the synthesis of mRNA degradation intermediates as the substrate for later cytoplasmic re-polyadenylation in both mouse and human (Fig. 7). Our recent studies reveal that Tut4/7 can incorporate U residues into maternal RNA, promoting maternal RNA degradation in mouse 2C embryos, and Tent4a/b can incorporate G residues into maternal mRNA, stabilizing them after cytoplasmic re-polyadenylation^3,5^. Interestingly, the length and non-A residues for poly(A) tails on the degradation intermediates are different from those on regular polyadenylation sites (Fig. 7). The function and mechanism of this difference await further investigation. Extensive cytoplasmic polyadenylation has also been seen on mRNA degradation intermediates in zebrafish^40^. Therefore, cytoplasmic polyadenylation on mRNA degradation intermediates is conserved in vertebrates, suggesting the functional importance of these re-polyadenylated mRNA degradation intermediates. However, the mechanism of production of the mRNA degradation intermediates in zebrafish remains to be explored. More broadly, it will be interesting to see whether global cytoplasmic polyadenylation on mRNA degradation intermediates and its production mechanism is conserved in OET of other model organisms, such as *Xenopus, Drosophila*, and *C. elegans*.

**Fig. 7.**
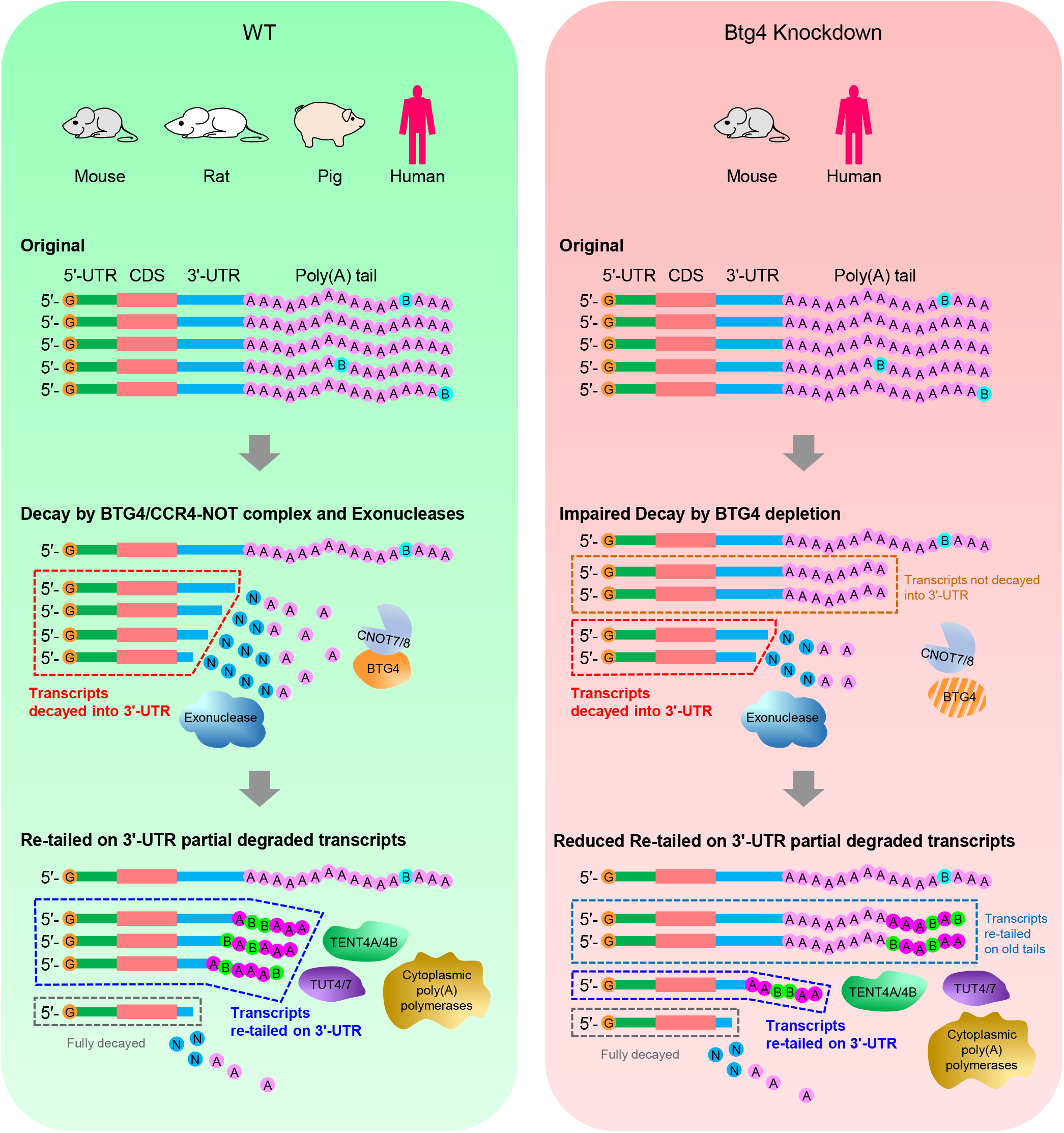
Model for the synthesis of non-classical poly(A) tails during mouse, rat, pig and human oocyte-to-embryo transition. Illustration of the generation of non-classical poly(A) tails during OET in mouse, rat, pig and human (left). The mRNA transcripts with classical 3′-UTR first experience global deadenylation in a Btg4 deadenylase-dependent manner to generate transcripts with a partially degraded 3′-UTR. Next, these transcripts with partially degraded 3′-UTR can be re-polyadenylated during OET to generate a non-classical poly(A) tail containing 3′-UTR partially degraded mRNA transcripts. If global deadenylation is blocked, for example by Btg4 depletion in mouse and human, the global deadenylation-mediated mRNA decay is impaired. Then, fewer non-classical poly(A) tails can be generated (right).

For the majority of eukaryotic mRNA, degradation starts from deadenylation of its poly(A) tail, followed by rapid degradation of the mRNA body by exonucleases^41^. Although a large fraction of maternal mRNA is fully degraded during oocyte maturation, a significant proportion of the maternal mRNA does not undergo full degradation but stabilized in the form of mRNA degradation intermediates. Among these mRNA degradation intermediates, a small fraction of them begins cytoplasmic polyadenylation at the MII stage, while the majority of them experience cytoplasmic polyadenylation after fertilization. This indicates that the deadenylation and full decay of mRNA are decoupled during the mammalian OET process, which is different from our previous knowledge in non-reproductive cells. Therefore, an interesting question remains to be answered as what mechanism protects the mRNA degradation intermediates from further degradation before cytoplasmic polyadenylation in the mammalian oocyte and embryos.

## Materials and Methods

### PAIso-seq1 and PAIso-seq2

The PAIso-seq1 and PAIso-seq2 libraries were constructed following the PAIso-seq1 and PAIso-seq2 protocols starting with purified total RNA as described previously^27,32^. The libraries were sequenced using PacBio Sequel I or Sequel II instruments at Annoroad.

### PAIso-seq1 and PAIso-seq2 data processing

The PAIso-seq1 and PAIso-seq2 data pre-processing, poly(A) tail sequence extraction, poly(A) tail length measurement and detection of non-A residues in poly(A) tails were conducted according to published procedures^5^.

### Polyadenylation site calling for annotated mRNA

For each species of the OET datasets, the GV stage data were used for polyadenylation site calling for annotated genes encoding mRNA. Clean CCS reads were aligned to the reference genomes using *minimap2*^42^. A greedy strategy was used for computing the depth of each candidate poly(A) site (namely the 3′-end of the alignment) as the number of reads aligned within a window of 5 nucleotides of the site. The site with maximum read depth across all candidates was added to the list of poly(A) sites if the site had a depth of at least 10 and did not occur within 20 nucleotides of a previously added site. The site was then removed from the list of candidates and the algorithm proceeded until no candidate sites remained. Protein-coding genes in the nuclear genome were included in this analysis.

### Classification of classical and non-classical polyadenylation sites

For each species of the OET datasets, the polyadenylation sites called in the GV stage data were used as reference point for classification of polyadenylation sites as classical or non-classical polyadenylation sites. Polyadenylation sites assigned to a given gene and located within 5 nt of the reference polyadenylation sites for the given gene were classified as classical polyadenylation sites. Polyadenylation sites assigned to a given gene and located 5 nt away from all reference polyadenylation sites for the given gene were classified as non-classical polyadenylation sites. Protein-coding genes in the nuclear genome were included in this analysis.

### Analysis of nucleotide composition around polyadenylation sites

Genomic sequences were extracted from upstream (−50 nt) and downstream (+50 nt) of the analyzed polyadenylation sites. Then, the base composition of each position was calculated using the *consensusMatrix* function in the *Biostrings* package and plotted using the *matplot* function in *R*. Protein-coding genes in the nuclear genome were included in this analysis. The presence of polyadenylation signal (PAS) sequence (AATAAA) and its most frequent variant (ATTAAA) were searched in the 51 nt sequence containing the upstream (−50 nt) to the polyadenylation sites for each transcript.

## Data Availability

The ccs data in bam format from PAIso-seq1and 2 experiments will be available at Genome Sequence Archive hosted by National Genomic Data Center. Custom scripts used for data analysis will be available upon request. This study includes analysis of the following published data: Legnini et al. (Gene Expression Omnibus database (GEO) accession no. GSE126465).

## Acknowledgements

This work was supported by the National Key Research and Development Program of China (2018YFA0107001, 2016YFA0100200), the Strategic Priority Research Program of the Chinese Academy of Sciences (XDA24020203), National Natural Science Foundation of China (31970588, 32170606), Natural Science Foundation of Heilongjiang province (YQ2020C003), the China Postdoctoral Science Foundation (2020M670516, 2020T130687), the State Key Laboratory of Molecular Developmental Biology, and Heilongjiang Touyan Innovation Team Program.

## Author Contributions

Yusheng Liu, Jiaqiang Wang and Falong Lu conceived the project and designed the study. Yusheng Liu, Yiwei Zhang, Hu Nie, Jiaqiang Wang and Falong Lu analyzed the sequencing data. Zhonghua Liu provided funding supports for the project. Yusheng Liu and Jiaqiang Wang organized all figures. Yusheng Liu, Jiaqiang Wang and Falong Lu supervised the project. Yusheng Liu, Jiaqiang Wang and Falong Lu wrote the manuscript with the input from the other authors.

## Figure legends

**Extended Data Fig. 1.**
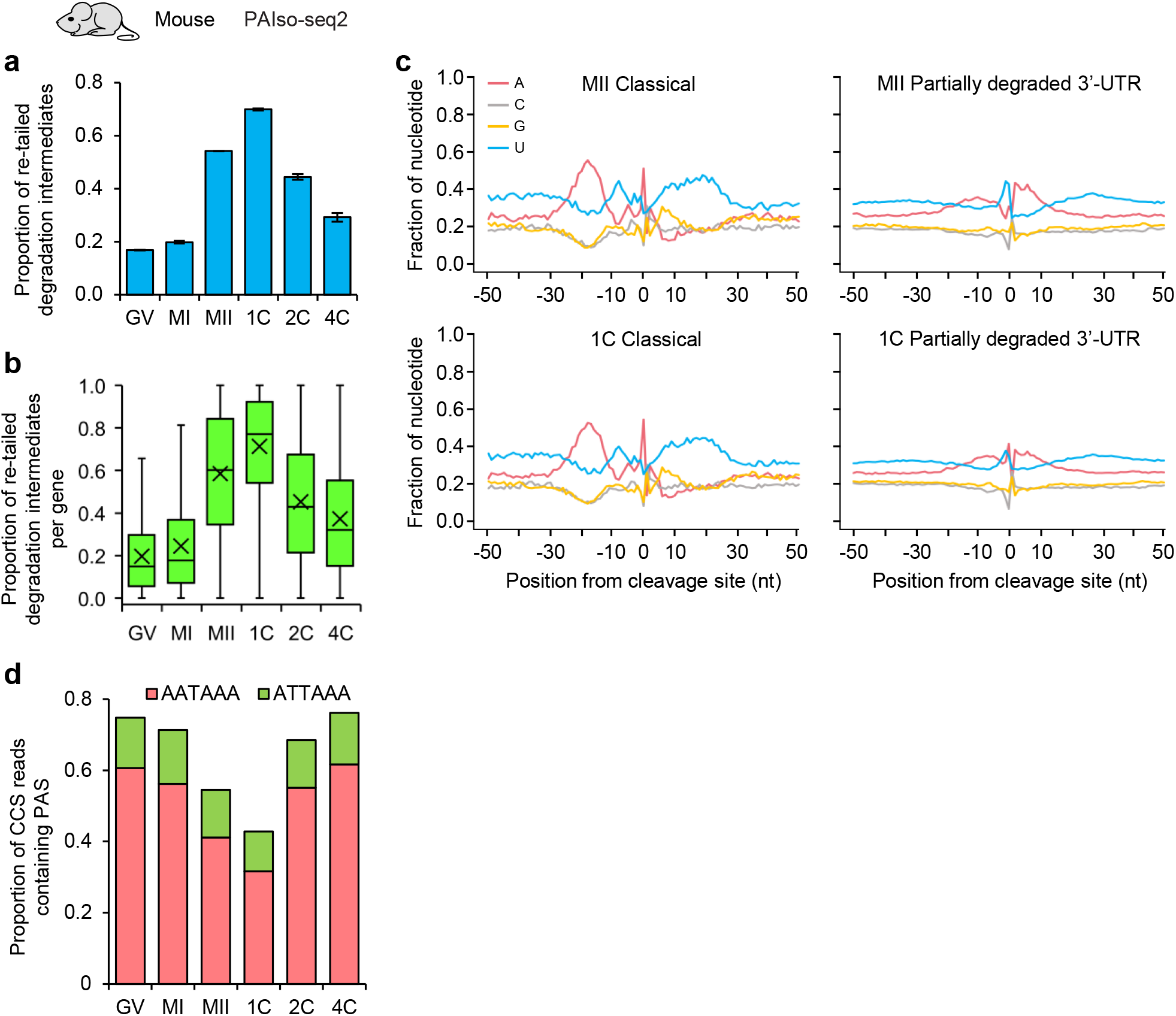
Re-polyadenylated mRNA degradation intermediates dominate polyadenylated mRNA during mouse oocyte-to-embryo transition revealed by PAIso-seq2. **a**, Overall proportion of the re-polyadenylated mRNA degradation intermediates for each of the 6 stages of mouse samples sequenced by PAIso-seq2. **b**, Box plot of the proportion of the re-polyadenylated mRNA degradation intermediates for each gene in each of the 6 stages of mouse samples sequenced by PAIso-seq2. Genes (n = 2,841) with at least 10 transcripts with a poly(A) tail of at least 1 nt in each of the 6 stages are included in the analysis. The “×” indicates the mean value, the black horizontal bars show the median value, and the top and bottom of the box represent the value of 25^th^ and 75^th^ percentile, respectively. **c**, Profile of nucleotide frequencies of the genomic sequences in the ± 50 nt vicinity of the last base before mRNA poly(A) tails for the combined transcripts with poly(A) tails from classical (left) and non-classical (right) polyadenylation sites in MII (top) and 1C (bottom) mouse samples sequenced by PAIso-seq2. **d**, Overall proportion of mRNA transcripts with canonical polyadenylation signal (PAS) in each of the 6 stage mouse samples sequenced by PAIso-seq2. Error bars indicate the SEM from two replicates (n=2). Transcripts with a poly(A) tail of at least 1 nt are included in the analysis.

**Extended Data Fig. 2.**
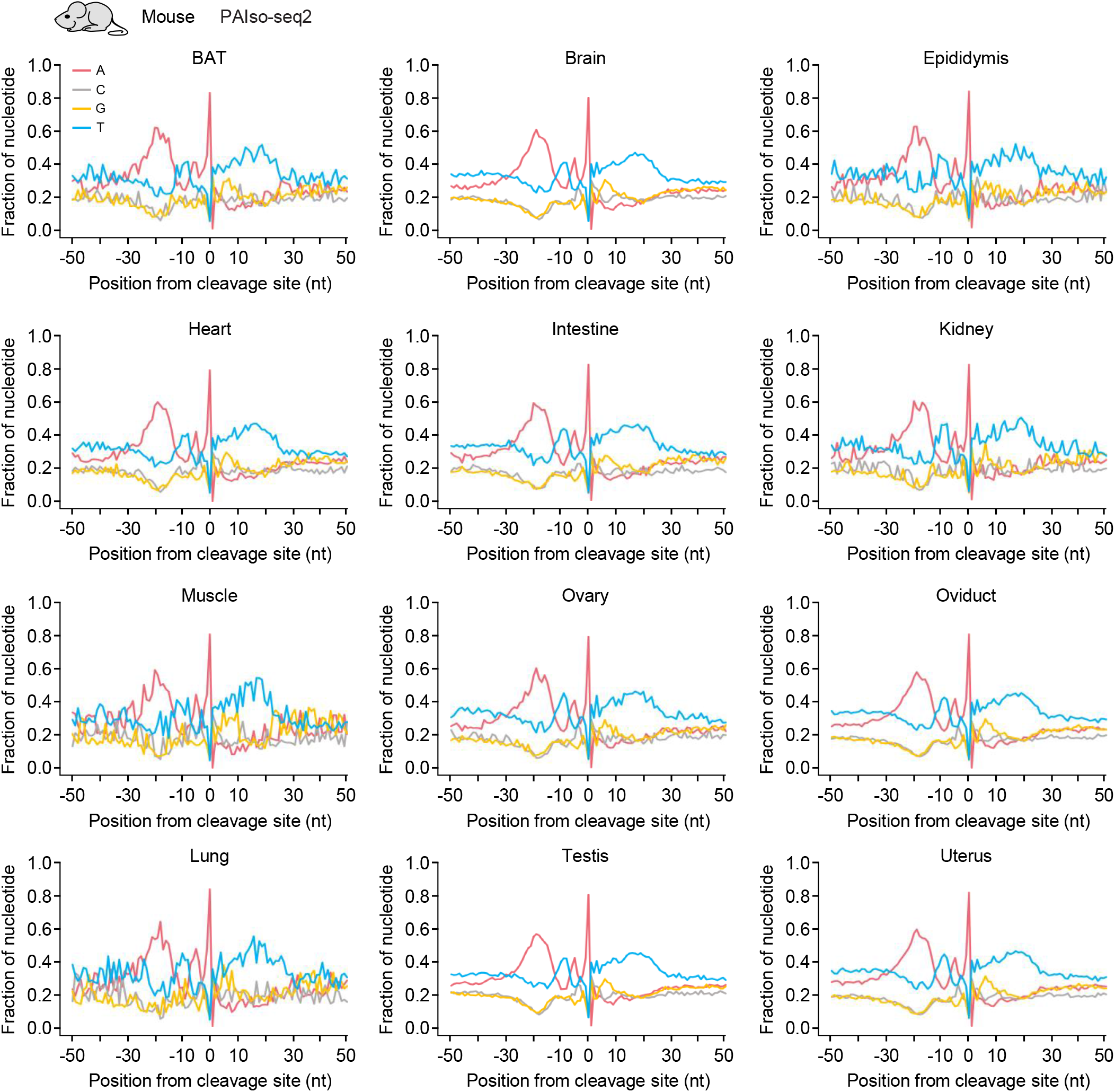
Similar nucleotide composition around the polyadenylation sites in somatic cells from different mouse tissues. Profile of nucleotide frequencies of the genomic sequences in the ± 50 nt vicinity of the last base before mRNA poly(A) tails sequenced by PAIso-seq2 in 12 different mouse tissues, including brown adipose tissue (BAT), brain, epididymis, heart, intestine, kidney, muscle, ovary, oviduct, lung, testis, and uterus.

**Extended Data Fig. 3.**
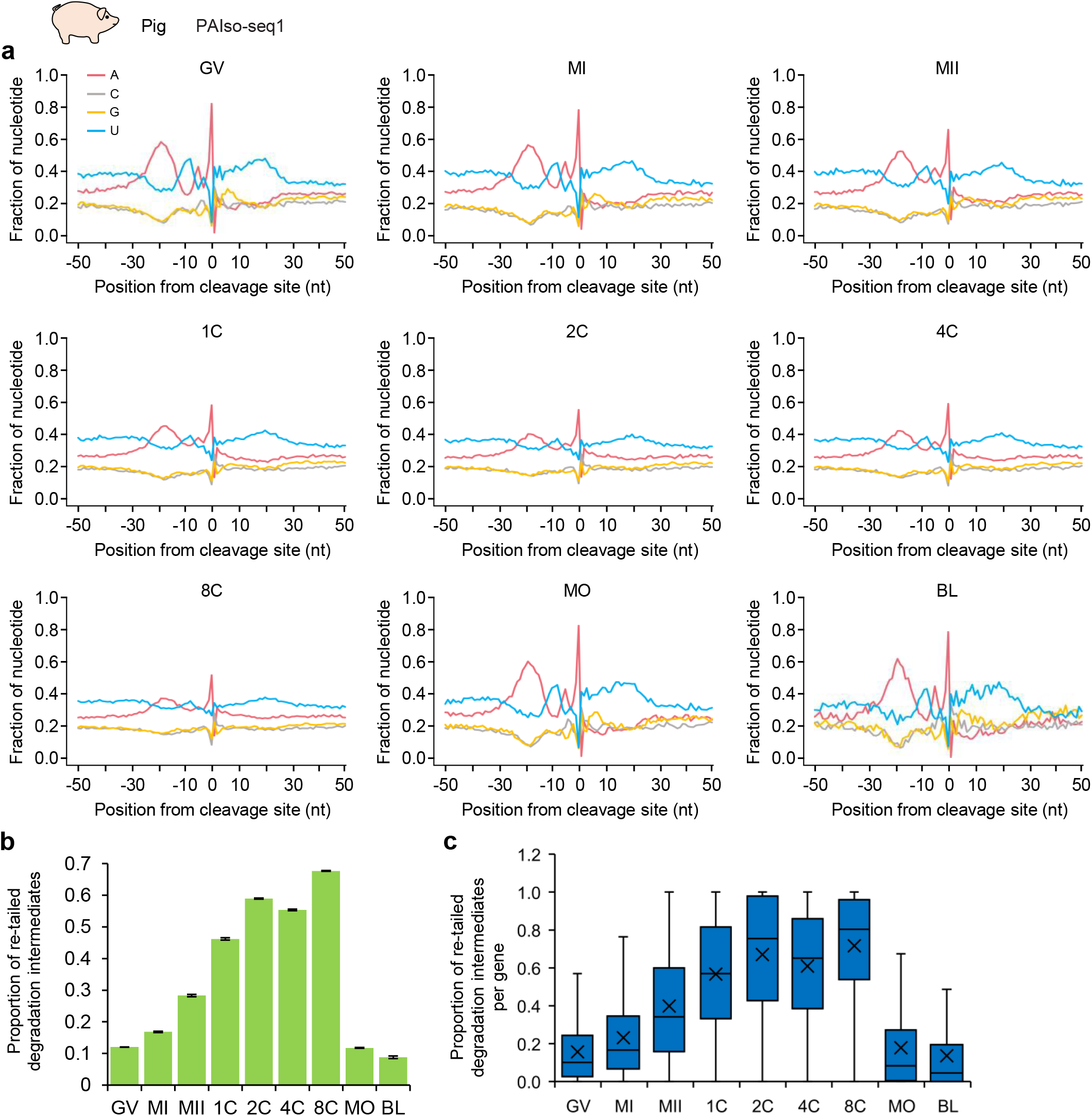
Re-polyadenylated mRNA degradation intermediates dominate polyadenylated mRNA during pig oocyte-to-embryo transition revealed by PAIso-seq1. **a**, Profile of nucleotide frequencies of the genomic sequences in the ± 50 nt vicinity of the last base before mRNA poly(A) tails sequenced by PAIso-seq1 in samples from different stages during pig OET, including GV, MI and MII stage oocytes, as well as 1C, 2C, 4C, 8C (8-cell), MO (morula) and BL (blastocyst) embryos. **b**, Overall proportion of the re-polyadenylated mRNA degradation intermediates for each of the 9 stages pig PAIso-seq1 samples. **c**, Box plot of the proportion of the re-polyadenylated mRNA degradation intermediates for each gene in each of the 9 stages pig PAIso-seq1 samples. Genes (n = 3,145) with at least 10 transcripts with poly(A) tail of at least 1 nt in each of the 9 stages are included in the analysis. Error bars indicate the SEM from two replicates (n=2). Transcripts with poly(A) tail of at least 1 nt are included in the analysis. For all the box plots, the “×” indicates the mean value, the black horizontal bars show the median value, and the top and bottom of the box represent the value of 25^th^ and 75^th^ percentile, respectively.

**Extended Data Fig. 4.**
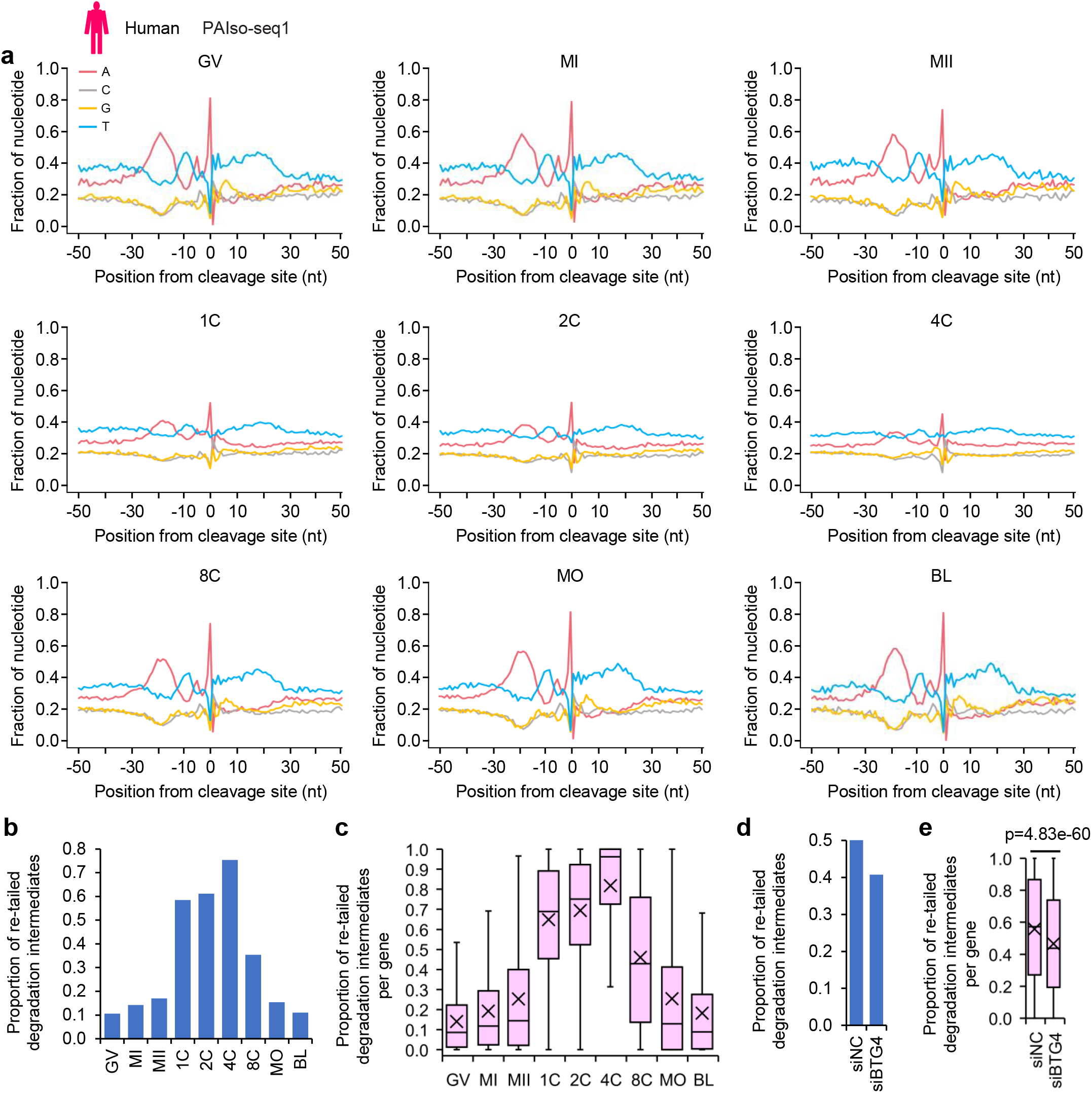
Re-polyadenylated mRNA degradation intermediates dominate polyadenylated mRNA during human oocyte-to-embryo transition revealed by PAIso-seq1. **a**, Profile of nucleotide frequencies of the genomic sequences in the ± 50 nt vicinity of the last base before mRNA poly(A) tails sequenced by PAIso-seq1 in samples from different stages during human OET, including GV, MI and MII stage oocytes, as well as 1C, 2C, 4C, 8C, MO and BL embryos. **b**, Overall proportion of the re-polyadenylated mRNA degradation intermediates for each of the 9 stages human PAIso-seq1 samples. **c**, Box plot of the proportion of the re-polyadenylated mRNA degradation intermediates for each gene in each of the 9 stages of human PAIso-seq1 samples. Genes (n = 1,577) with at least 10 transcripts with poly(A) tail of at least 1 nt in each of the 9 stages are included in the analysis. **d**, Overall proportion of the re-polyadenylated mRNA degradation intermediates in siNC and siBTG4 human 1C embryos sequenced by PAIso-seq1. **e**, Box plot of the proportion of the re-polyadenylated mRNA degradation intermediates for each gene in siNC and siBTG4 human 1C embryos sequenced by PAIso-seq1. The *p* value is calculated by Student’s *t*-test. Genes (n = 5,824) with at least 10 transcripts with a poly(A) tail of at least 1 nt in both samples are included in the analysis. Transcripts with a poly(A) tail of at least 1 nt are included in the analysis. For all box plots, the “×” indicates the mean value, the black horizontal bars show the median value, and the top and bottom of the box represent the value of 25^th^ and 75^th^ percentile, respectively.

